# Marked enhancement of neutralizing antibody and IFN-γ T-cell responses by GX-19N DNA booster in mice primed with inactivated vaccine

**DOI:** 10.1101/2021.11.02.467026

**Authors:** Yong Bok Seo, Duckhyang Shin, You Suk Suh, Juyoung Na, Ji In Ryu, Young Chul Sung

## Abstract

In response to the COVID-19 pandemic, an unprecedented level of vaccine development has occurred. As a result, various COVID-19 vaccines have been approved for use. Among these, inactivated virus particle (VP) vaccines have been widely used worldwide, but additional vaccination strategies are needed because of the short duration of immune responses elicited by these vaccines. Here, we evaluated homologous and heterologous prime–boost regimens using a VP vaccine and GX-19N DNA vaccine for their ability to enhance the protective immune response against SARS-CoV-2. We demonstrated that a heterologous prime–boost regimen with the VP vaccine and GX-19N DNA vaccine resulted in enhanced S_RBD_- & N-specific antibody responses, compared to the homologous VP vaccine prime–boost vaccination. In addition, the neutralizing antibody response was significantly improved with the heterologous VP prime–DNA boost regimen, and the neutralizing antibody induced with the heterologous prime–boost regimen did not decrease against the SARS-CoV-2 variant of concern (VOC). The heterologous VP prime–DNA boost regimen not only significantly increased S- and N-specific IFN-γ T-cell responses, but also induced an equivalent level of T-cell response against SARS-CoV-2 VOCs. Our results provide new insights into prophylactic vaccination strategies for COVID-19 vaccination.

## MAIN TEXT

There has been an unprecedented level of vaccine development in response to the COVID-19 pandemic. As a result, 22 COVID-19 vaccines have been approved for use, and more than 3.7 billion doses have been administered. This rapid development of an effective vaccine against COVID-19 and its distribution to the general population has proven to be a very successful strategy for reducing the transmission and disease burden of COVID-19.

Among several vaccine platforms, inactivated virus particle (VP) vaccines have been extensively studied. They are generally safe and are widely used to prevent respiratory infections such as influenza and other infectious diseases such as hepatitis A, polio, and rabies^1^. VP vaccines are manufactured through an inactivation process of the virus, which leads to a loss of infectivity but retains the main viral antigenicity. VP vaccines also have the advantage of being easily stored and transported for years at 2–8 °C, making them suitable for use in many low-income countries and locations with limited cold storage capacity. However, production rates may be limited by the yield of the virus in cell culture and the requirements for a production facility at biosafety level 3. The COVID-19 VP vaccines BBIBP-CorV (Sinopharm Beijing), Coronavac (Sinovac), and BBV152 (Bharat Biotech) have demonstrated protective efficacies of 78.1%^2^, 50.7%^3^, and 77.8%^4^, respectively, and have been approved for emergency use in several countries. As a result, a significant portion of the world’s population has received a two-dose emergency immunization with one of the COVID-19 VP vaccines.

The DNA vaccine is an effective platform for rapid control of the SARS-CoV-2 outbreak as it can be completed in weeks to produce a clinical-grade vaccine^5^. In addition, DNA vaccines have the advantage of being highly stable at various temperatures, making them suitable for delivery in environments where resources are limited and there are no cold chains^6^. It has been reported that unlike conventional vaccines, DNA vaccines can effectively stimulate both cellular and humoral responses to pathogens in a challenge model^7^. In addition, the potential of prophylactic DNA vaccines against viral infections such as MERS-CoV and ZIKV has been demonstrated in recent clinical trials^8, 9, 10, 11^. As the COVID-19 pandemic has spread worldwide, recent studies have reported that DNA vaccines induce antigen-specific T-cell responses, neutralize antibodies, and further protect animals from SARS-CoV-2 challenge^12, 13, 14^. A COVID-19 DNA vaccine, ZyCoV-D, has recently been approved for emergency use as it has shown 66.6% protective efficacy in phase 3 clinical trials^15^. In addition, two, eight, and one candidate DNA vaccines are currently in phases 3, 2, and 1 of clinical trials, respectively^16^.

A heterologous prime–boost has been shown to effectively improve the immunogenicity of HIV-1 and influenza vaccines^17, 18, 19, 20^. Heterologous prime-boost regimens using DNA and VP vaccines have been shown to induce more effective antibody and T-cell responses than the homologous prime-boost regimen^16, 21^. Although most previous studies have used heterologous DNA vaccine prime–VP vaccine boost regimens, given the current situation where COVID-19 VP vaccines are predominant and COVID-19 DNA vaccines are rare, efficacy evaluation of heterogeneous VP vaccine prime–DNA vaccine boost regimens could provide more practical and valuable information.

In this study, six-week-old female mice were primed with a VP vaccine and boosted with the VP vaccine or GX-19N DNA vaccine four weeks later for booster vaccination. Immune responses were evaluated two weeks after the last vaccination. The Spike-specific binding antibody was evaluated by ELISA using the receptor binding domain (S_RBD_, Wuhan) and nucleocapsid (N, Wuhan) proteins. The heterologous VP vaccine prime–GX-19N DNA vaccine boost induced a significantly higher S_RBD_-specific antibody response than the homologous VP vaccine prime–boost regimen. Compared to VP prime, VP boost increased antibody titer by 1.7-fold, whereas DNA boost increased antibody titer by 181-fold. (Fig. 1A). The ratio of IgG2a to IgG1 antibody titer was, albeit statistically insignificant, increased after GX-19N boosting (Fig 1B), indicating the enhancement of Th1-polarized immunity, as reported previously^22, 23^. Likewise, the heterologous prime-boost vaccination also induced significantly higher NP-specific antibody responses than the homologous prime-boost vaccination. VP-prime followed by VP-boost increased antibody titer by 1.4-fold, whereas GX-19N DNA-boost increased antibody titer by 512-fold. (Fig. 1C). In addition, we evaluated the neutralizing antibody responses to the Wuhan and VOCs (B.1.351 and B.1.617.2) using the surrogate virus neutralization test (sVNT), which is highly correlated with the conventional virus neutralization test (cVNT) and pseudovirus-based VNT (pVNT)^24^. In contrast to the previous report that the boosting with 2 μg VP vaccine increased about 2-fold neutralization titer, the booster immunization with VP vaccine in our experiment did not further enhance neutralization antibody responses in mice primed with VP vaccine. This discrepancy may be due to the differences in immunization dose (0.4 μg vs. 2 μg), and assay methods for evaluating neutralization antibody responses (sVNT vs. cVNT).

**Figure 1.**
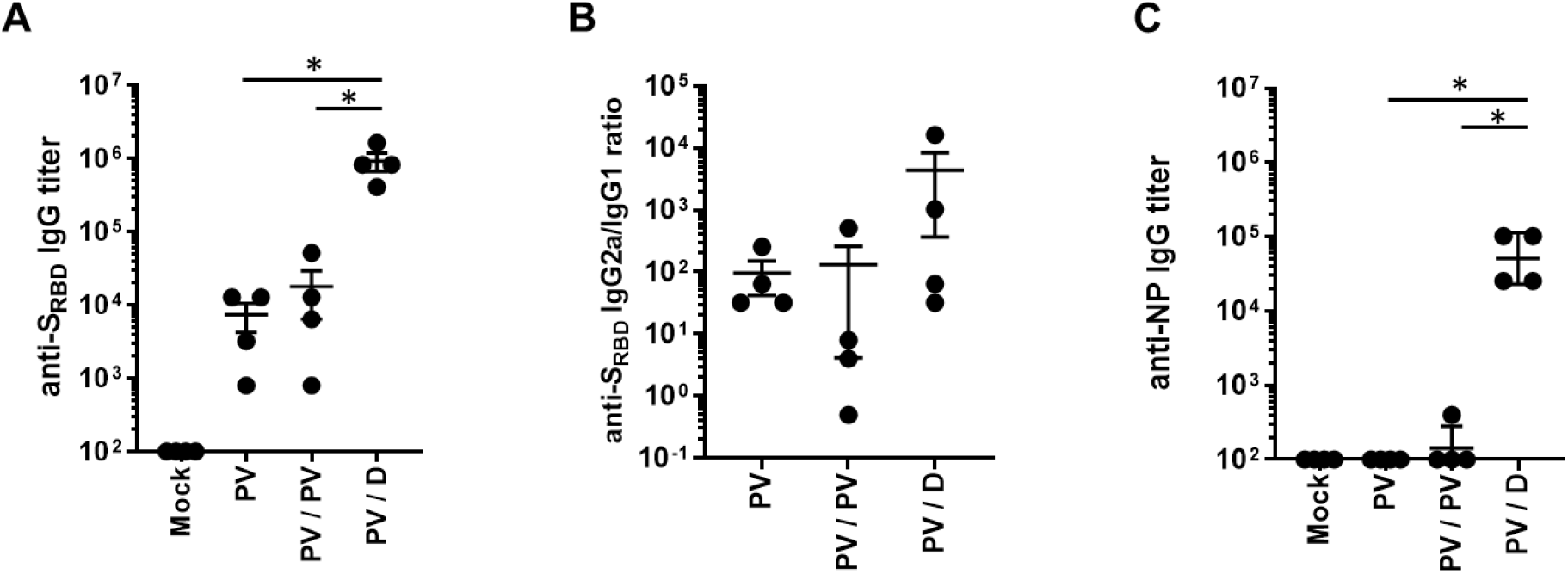
Heterologous VP prime–GX-19N DNA boost showed higher IgG titer than the homologous VP prime–boost regimen. Balb/c mice (n=4/group) were immunized at weeks 0 and 4 as described in the methods, and antibody responses were measured in the sera of mice collected 2 weeks after the last immunization. Graphs show SARS-CoV-2 S_RBD_-specific IgG titer (A), ratios of S_RBD_-specific IgG2a to IgG1 titer (B) and SARS-CoV-2 N-specific IgG titer (C). Individual mice were represented by a single data point. *P-values* were determined by two-tailed Student’s *t*-test. *p<0.05.

Heterologous vaccination induced a significantly higher neutralizing antibody titer than the homologous VP prime-boost vaccination. The neutralizing antibody titer obtained through the heterogeneous regimen was 76.1-fold higher for the Wuhan (mean = 14 vs. 1,076 sVNT_20_ titer), 53.8-fold higher for the B.1.351 (mean = 10 vs. 538 sVNT_20_ titer), and 76.1-fold higher for the B.1.617.2 (mean = 14 vs. 1,076 sVNT_20_ titer) variants than that obtained through the homologous prime-boost regimen (Fig 2A). Interestingly, the neutralizing antibody titer obtained for the B.1.351 and B.1.617.2 variants through the heterologous regimen were comparable to those obtained for the Wuhan (Fig. 2B), presumably due to the induction of cross-reactive neutralizing antibodies. It is worth noting that level of neutralization antibody responses induced by an mRNA vaccine (BNT162b2) was reduced about 5-fold against variants (B.1.351, B.1.617.2), compared to the wild-type^25^.

**Figure 2.**
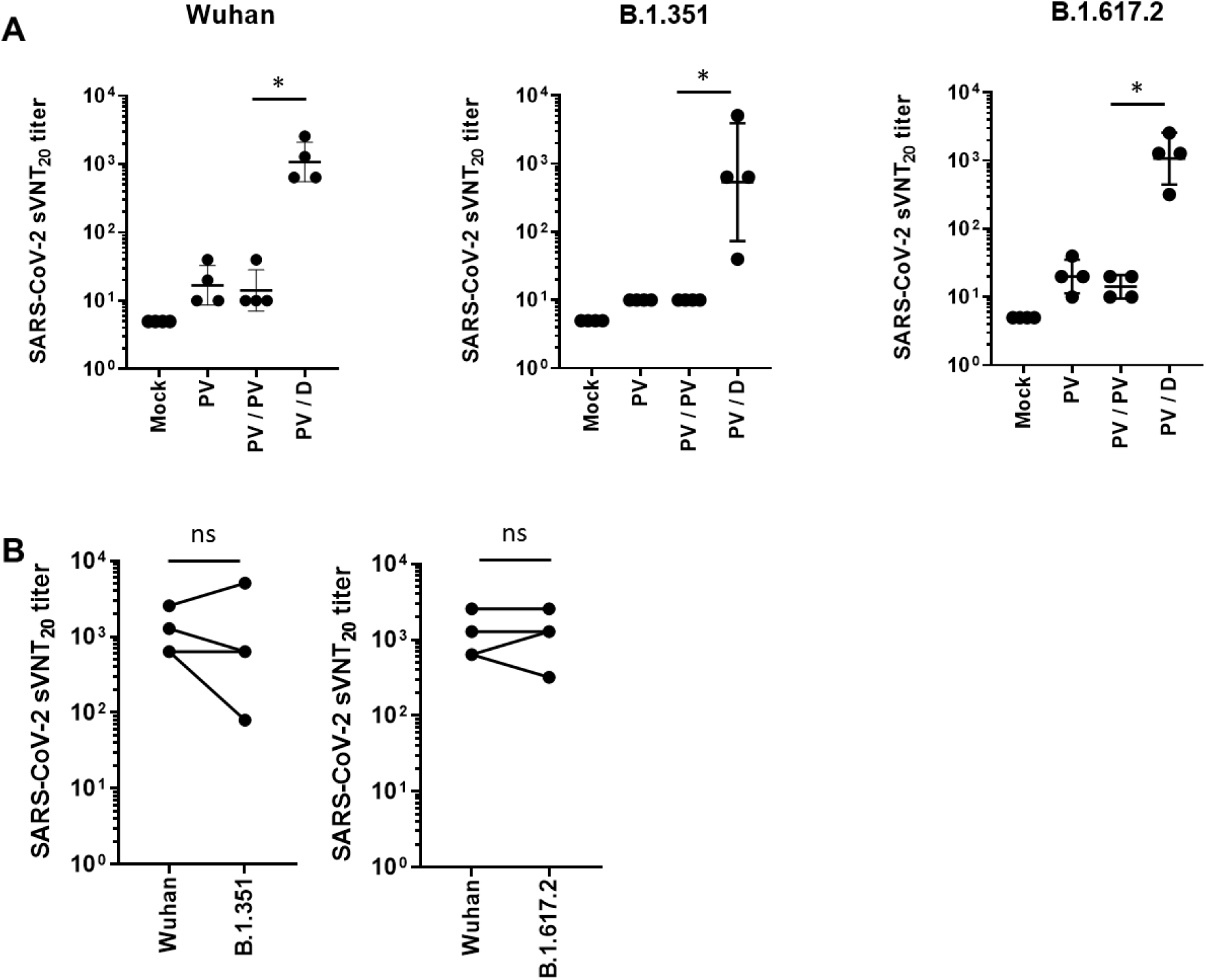
Surrogate virus neutralization titers against SARS-CoV-2 Wuhan, B.1.351, and B.1.617.2. Balb/c mice (n=4/group) were immunized at weeks 0 and 4 as described in the methods, and antibody responses were measured in the sera of mice collected 2 weeks after the last immunization. Sera from vaccinated mice were tested for sVNT_20_ titers against Wuhan, B.1.351, and B.1.617.2 (A). sVNT_20_ titer against Wuhan compared to that against B.1.351 and B.1.617.2 (B). Individual mice are represented by a single data point. *P-values* were determined by two-tailed Student’s *t*-test. ns, not significant; *p<0.05.

T-cell responses were also evaluated following the homologous and heterologous vaccination regimens. A higher Wuhan SARS-CoV-2 spike-specific IFN-γ response was detected by ELIspot in mice that received a heterologous vaccination regimen. Compared to VP-prime, VP-boost increased T cell response by 1.6-fold, whereas GX-19N DNAboost increased T cell response by 38.3-fold. (Fig. 3A). As expected, we observed similar levels of cellular responses to the B.1.351 (mean = 1,644 SFUs/10^6^ splenocytes) and B.1.617.2 (mean = 1,753 SFUs/10^6^ splenocytes) spike peptides (Fig. 3B). Similar to the results for the spike peptides, a higher N-specific IFN-γ response was observed in the heterologous VP vaccine prime–GX-19N DNA vaccine boost regimen. VP-prime followed by VP-boost increased cellular response by 2.9-fold, whereas GX-19N DNA-boost increased cellular response by 8.8-fold. (Fig. 4A). We also observed similar levels of IFN-γ T cell response to B.1.351 (mean = 739 SFUs/10^6^ splenocytes) and B.1.617.2 (mean = 794 SFUs/10^6^ splenocytes) nucleocapsid peptides (Fig. 4B). This is consistent with previous findings that cellular immunity is relatively unimpaired by VOCs compared to neutralizing antibody responses ^26^. Similar results showing enhancement of neutralization antibody and T cell responses by GX-19N booster immunization were also obtained in a separate independent experiments with different dose and boosting interval.

**Figure 3.**
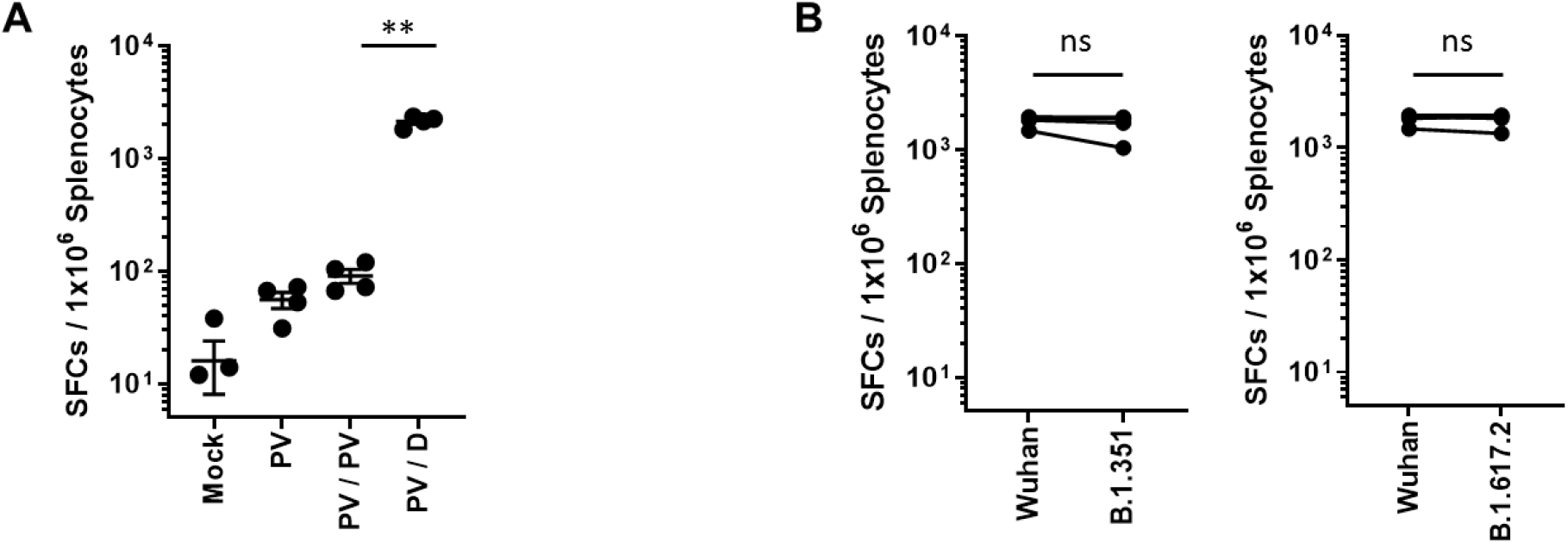
Spike-specific T-cell response against SARS-CoV-2 Wuhan, B.1.351, and B.1.617.2. Balb/c mice (n=4/group) were immunized at weeks 0 and 4 as described in the methods, and the T-cell response was measured by IFN-γ ELIspot in splenocytes stimulated with peptide pools spanning the SARS-CoV-2 spike protein. Spike-specific T cells (A). T-cell response against Wuhan compared to that against B.1.351 and B.1.6172. (B). Individual mice are represented by a single data point. *P*-values were determined by two-tailed Student’s *t*-test. ns, not significant; **p<0.01.

**Figure 4.**
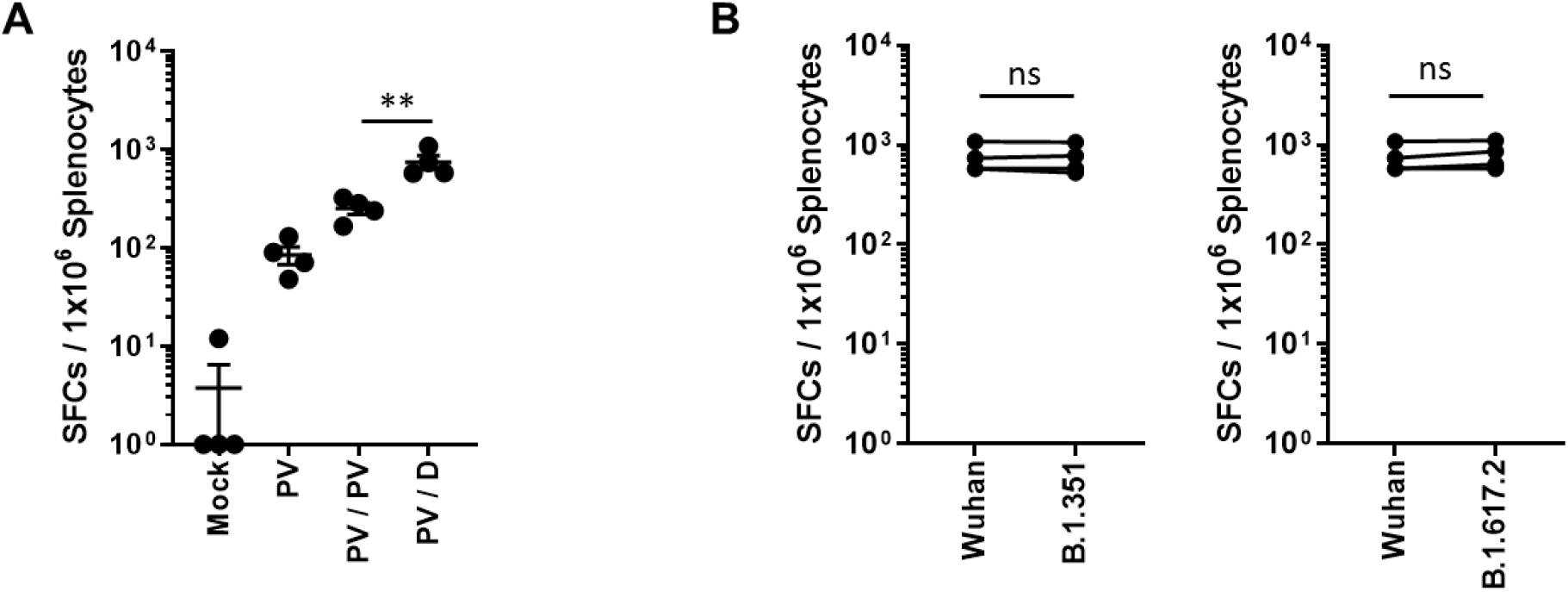
Nucleocapsid-specific T-cell response against SARS-CoV-2 Wuhan, B.1.351, and B.1.617.2. Balb/c mice (n=4/group) were immunized at weeks 0 and 4 as described in the methods, and the T-cell response was measured by IFN-γ ELIspot in splenocytes stimulated with peptide pools spanning the SARS-CoV-2 nucleocapsid protein. Nucleocapsid-specific T-cells (A). T-cell response against Wuhan compared to that against B.1.351 and B.1.6172 (B). Individual mice are represented by a single data point. *P*-values were determined by two-tailed Student’s *t*-test. ns, not significant; **p<0.01.

Clinical studies are underway to compare the immunogenicity of heterologous prime–boost regimens using approved COVID-19 vaccines. Some studies have already reported that heterologous prime–boost immunization induces a superior immune response compared to homologous prime– boost immunization^27, 28, 29^. However, these studies are limited to adenoviral vector-based vaccines (ChAdOx1nCoV-19) and mRNA-based vaccines (BNT162b2), and heterologous prime–boost studies for other platform vaccines such as DNA vaccines and VP vaccines are not in progress. The COVID-19 VP vaccine is one of the most commonly used vaccines in the world. However, some studies have demonstrated that a significant proportion of antibody responses are reduced within six months of vaccination, indicating the need for an additional immunization strategy for VP vaccines^30, 31, 32^, and one plausible approach may be a heterologous prime–boost immunization strategy. Here, we have shown that a highly effective antibody response and T-cell response can be induced by heterologous prime–boost using the VP vaccine and the GX-19N DNA vaccine. Thus, GX-19N boost vaccination is likely to compensate for the weakness of VP vaccines, which are known to induce weak T-cell responses compared to other vaccine platforms such as DNA, mRNA, and viral vectors ^33, 34, 35, 36^. Finally, based on our results, together with good safety and stability of DNA vaccine, GX-19N DNA vaccine is expected to be optimal as a heterologous booster for VP vaccines.

## Methods

### Vaccines

The COVID-19 DNA vaccine, consisting of GX-19 and GX-21 at a 1:2 ratio, was constructed by inserting the antigen genes of SARS-CoV-2 into the pGX27 vector. GX-19 (pGX27-S_ΔTM/IC_) contains the SARS-CoV-2 spike (S) gene lacking the transmembrane (TM)/intracellular (IC) domain, and GX-21 (pGX27-S_RBD_-F/NP) is designed to express the fusion protein of the receptor binding domain (RBD) of the spike protein, the T4 fibritin C-terminal foldon (S_RBD_-Foldon), and the nucleocapsid protein (N). The S, S_RBD_-Foldon, and N are preceded by the secretory signal sequence of the tissue plasminogen activation (tPA). The inactivated SARS-CoV-2 vaccine produced from Vero cells contains 4 μg of viral antigens and 0.225 mg of aluminum hydroxide adjuvant in a 0.5-mL dose.

### Mouse immunizations

Female BALB/c mice aged 6–8 weeks (Central Lab Animal) were intramuscularly immunized with 0.4 μg/ animal VP vaccine at week 0 and boosted with 12 μg/animal GX-19N, in a total volume of 50 μL PBS, into the tibialis anterior muscle with *in vivo* electroporation with the OrbiJector® system (SL VAXiGEN Inc.) at week 4. The mice were sacrificed two weeks after the final immunization.

### Antigen binding ELISA

Serum collected at each time point was evaluated for binding titers. In this assay, 96-well immunosorbent plates (NUNC) were coated with 1 μg/mL recombinant SARS-CoV-2 Spike RBD-His protein (Sino Biological 40592-V08B) in PBS overnight at 4 °C. Plates were washed three times with 0.05% PBST (Tween 20 in PBS) and blocked with 5% skim milk in 0.05% PBST (SM) for 2–3 h at room temperature. Sera were serially diluted in 5% SM, added to the wells, and incubated for 2 h at 37 °C. Following incubation, plates were washed five times with 0.05% PBST and then incubated with horseradish peroxidase (HRP)-conjugated anti-mouse IgG (Jackson ImmunoResearch Laboratories 115-035-003), IgG1 (Jackson ImmunoResearch Laboratories 115-035-205), or IgG2a (Jackson ImmunoResearch Laboratories 115-035-206) for 1 h at 37 °C. After the final wash, plates were developed using TMB solution (Surmodics TMBW-1000-01), and the reaction was stopped with 2N H_2_SO_4_. The plates were read at 450 nm using a SpectraMax Plus384 (Molecular Devices).

### Surrogate virus-neutralization assay

Surrogate virus neutralization test (sVNT) analyzed the binding ability of RBD to ACE2 after neutralizing RBD with antibodies in serum. Serum collected two weeks after final immunization was quantified according to the manufacturer’s instructions (Sugentech, CONE001E). Briefly, sera were serially diluted in dilution buffer and reacted with HRP-conjugated RBD for 30 min at 37 °C. The reacted samples were added to a plate coated with human ACE2 protein and incubated for 15 min at 37 °C. Following incubation, the plates were washed five times with wash solution. After the final wash, plates were developed using TMB solution, the reaction was arrested with a stop solution. The plates were read at 450 nm using a SpectraMax Plus384 (Molecular Devices). The reciprocal of the dilution resulting in a binding inhibition rate of 20% or more (PI20) was defined as the neutralizing antibody titer.

### IFN-γ ELISPOT

The Mouse IFN-γ ELISPOT set (BD 551083) was used as directed by the manufacturer. ELISPOT plates were coated with purified anti-mouse IFN-γ capture antibody and incubated overnight at 4 °C. Plates were washed and blocked for 2 h with RPMI + 10% FBS (R10 media) Then, 5 × 10^5^ splenocytes were added to each well and stimulated for 24 h at 37 °C in 5% CO_2_ with R10 media (negative control), concanavalin A (positive control), or specific peptide pools (2 μg/ml). Peptide pools consisted of 15-mer peptides overlapping by 11 amino acids and spanned the entire S and N proteins of SARS-CoV-2 (GenScript). After stimulation, the plates were washed, and spots were developed according to the manufacturer’s instructions. Plates were scanned and counted using the AID ELISPOT reader classic. Spot-forming units (SFU) per million cells was calculated by subtracting the number of negative control wells.

### Statistical analysis

Data analyses were performed using GraphPad Prism 7 (GraphPad Software). Comparison of data between groups was performed using two-tailed Student’s *t*-tests. Statistical significance was set at P <0.05.

## References

1. Kyriakidis, N.C., Lopez-Cortes, A., Gonzalez, E.V., Grimaldos, A.B. & Prado, E.O. SARS-CoV-2 vaccines strategies: a comprehensive review of phase 3 candidates. NPJ Vaccines 6, 28 (2021).

2. Al Kaabi, N. et al. Effect of 2 Inactivated SARS-CoV-2 Vaccines on Symptomatic COVID-19 Infection in Adults: A Randomized Clinical Trial. JAMA 326, 35–45 (2021).

3. Ricardo Palacios et al. Efficacy and safety of a COVID-19 inactivated vaccine in healthcare professionals in Brazil: The PROFISCOV study. SSRN, 2021

4. Ella, R. et al. Efficacy, safety, and lot to lot immunogenicity of an inactivated SARS-CoV-2 vaccine BBV152): a, double-blind, randomised, controlled phase 3 trial. MedRxiv, (2021).

5. Graham, B.S., Mascola, J.R. & Fauci, A.S. Novel Vaccine Technologies: Essential Components of an Adequate Response to Emerging Viral Diseases. JAMA 319, 1431–1432 (2018).

6. Tebas, P. et al. Intradermal SynCon(R) Ebola GP DNA Vaccine Is Temperature Stable and Safely Demonstrates Cellular and Humoral Immunogenicity Advantages in Healthy Volunteers. J Infect Dis 220, 400–410 (2019).

7. Alexander N. Zakhartchouk et al. Optimization of a DNA Vaccine Against SARS. DNA AND CELL BIOLOGY, 2007

8. Modjarrad, K. et al. Safety and immunogenicity of an anti-Middle East respiratory syndrome coronavirus DNA vaccine: a phase 1, open-label, single-arm, dose-escalation trial. The Lancet Infectious Diseases 19, 1013–1022 (2019).

9. Gaudinski, M.R. et al. Safety, tolerability, and immunogenicity of two Zika virus DNA vaccine candidates in healthy adults: randomised, open-label, phase 1 clinical trials. The Lancet 391, 552–562 (2018).

10. Tebas, P. et al. Safety and Immunogenicity of an Anti–Zika Virus DNA Vaccine. New England Journal of Medicine 385, e35 (2021).

11. Gary, E.N. & Weiner, D.B. DNA vaccines: prime time is now. Curr Opin Immunol 65, 21–27 (2020).

12. Smith, T.R.F. et al. Immunogenicity of a DNA vaccine candidate for COVID-19. Nat Commun 11, 2601 (2020).

13. Jingyou Yu et al. YDNA vaccine protection against SARS-CoV-2 in rhesus macaques. Science, 2020.

14. Seo, Y.B. et al. Soluble Spike DNA Vaccine Provides Long-Term Protective Immunity against SARS-CoV-2 in Mice and Nonhuman Primates. Vaccines (Basel) 9(2021).

15. https://www.zyduscadila.com/public/pdf/pressrelease/ZyCoV_D_Press_Release_1_7_2021.pdf

16. SUBHABRATA Biswas et al. Preexposure Efficacy of a Novel Combination DNA and Inactivated Rabies Virus Vaccine. HUMAN GENE THERAPY, 2001.

17. Wang, S. et al. Enhanced immunogenicity of gp120 protein when combined with recombinant DNA priming to generate antibodies that neutralize the JR-FL primary isolate of human immunodeficiency virus type 1. J Virol 79, 7933–7937 (2005).

18. Wang, S. et al. Polyvalent HIV-1 Env vaccine formulations delivered by the DNA priming plus protein boosting approach are effective in generating neutralizing antibodies against primary human immunodeficiency virus type 1 isolates from subtypes A, B, C, D and E. Virology 350, 34–47 (2006).

19. Bansal, A. et al. Multifunctional T-cell characteristics induced by a polyvalent DNA prime/protein boost human immunodeficiency virus type 1 vaccine regimen given to healthy adults are dependent on the route and dose of administration. J Virol 82, 6458–6469 (2008).

20. John W. Shiver et al.. Replication-incompetent adenoviral vaccine vector elicits effective antiimmunode®ciency-virus immunity. Nature, 2002.

21. Wang, S. et al. Heterologous HA DNA vaccine prime--inactivated influenza vaccine boost is more effective than using DNA or inactivated vaccine alone in eliciting antibody responses against H1 or H3 serotype influenza viruses. Vaccine 26, 3626–3633 (2008).

22. YS Lim et al.. Potentiation of antigen-specific, Th1 immune responses by multiple DNA vaccination with an ovalbumin/interferon-c hybrid construct. Immunology, 1998.

23. MK Song et al. Enhancement of Immunoglobulin G2a and Cytotoxic T-Lymphocyte Responses by a Booster Immunization with Recombinant Hepatitis C Virus E2 Protein in E2 DNA-Primed Mice. JOURNAL OF VIROLOGY, 2000.

24. Tan, C.W. et al. A SARS-CoV-2 surrogate virus neutralization test based on antibody-mediated blockage of ACE2-spike protein-protein interaction. Nat Biotechnol 38, 1073–1078 (2020).

25. Wall, E.C. et al. Neutralising antibody activity against SARS-CoV-2 VOCs B.1.617.2 and B.1.351 by BNT162b2 vaccination. The Lancet 397, 2331–2333 (2021).

26. Tarke, A. et al. Impact of SARS-CoV-2 variants on the total CD4(+) and CD8(+) T cell reactivity in infected or vaccinated individuals. Cell Rep Med 2, 100355 (2021).

27. Barros-Martins, J. et al. Immune responses against SARS-CoV-2 variants after heterologous and homologous ChAdOx1 nCoV-19/BNT162b2 vaccination. Nat Med 27, 1525–1529 (2021).

28. David Hillus et al. Safety, reactogenicity, and immunogenicity of homologous and heterologous prime-boost immunisation with ChAdOx1 nCoV-19 and BNT162b2: a prospective cohort study. Lancet Respir Med, 2021.

29. Liu, X. et al. Safety and immunogenicity of heterologous versus homologous prime-boost schedules with an adenoviral vectored and mRNA COVID-19 vaccine (Com-COV): a single-blind, randomised, non-inferiority trial. The Lancet 398, 856–869 (2021).

30. Kara, E. et al. Humoral immune response in inactivated SARS-CoV-2 vaccine: When should a booster dose be administered? MedRxiv, (2021).

31. Li, M. et al. A booster dose is immunogenic and will be needed for older adults who have completed two doses vaccination with CoronaVac: a randomised, double-blind, placebo-controlled, phase 1/2 clinical trial. MedRxiv, (2021).

32. Pan, H. et al. Immunogenicity and safety of a third dose, and immune persistence of CoronaVac vaccine in healthy adults aged 18-59 years: interim results from a double-blind, randomized, placebo-controlled phase 2 clinical trial. MedRxiv, (2021).

33. Xia, S. et al. Safety and immunogenicity of an inactivated SARS-CoV-2 vaccine, BBIBP-CorV: a randomised, double-blind, placebo-controlled, phase 1/2 trial. The Lancet Infectious Diseases 21, 39–51 (2021).

34. Xia, S. et al. Effect of an Inactivated Vaccine Against SARS-CoV-2 on Safety and Immunogenicity Outcomes: Interim Analysis of 2 Randomized Clinical Trials. JAMA 324, 951–960 (2020).

35. Zhang, Y. et al. Safety, tolerability, and immunogenicity of an inactivated SARS-CoV-2 vaccine in healthy adults aged 18–59 years: a randomised, double-blind, placebo-controlled, phase 1/2 clinical trial. The Lancet Infectious Diseases 21, 181–192 (2021).

36. Yang, S. et al. Safety and immunogenicity of a recombinant tandem-repeat dimeric RBD-based protein subunit vaccine (ZF2001) against COVID-19 in adults: two randomised, double-blind, placebo-controlled, phase 1 and 2 trials. The Lancet Infectious Diseases 21, 1107–1119 (2021).

